# ConsAlign: simultaneous RNA structural aligner based on rich transfer learning and thermodynamic ensemble model of alignment scoring

**DOI:** 10.1101/2022.04.27.489566

**Authors:** Masaki Tagashira

**Affiliations:** Department of Computational Biology and Medical Sciences, University of Tokyo, Chiba 277-8561, Japan

## Abstract

**Motivation:** To capture structural homology in RNAs, predicting RNA structural alignments has been a fundamental framework around RNA science. Learning simultaneous RNA structural alignments in their rich scoring is an undeveloped subject because evaluating them is computationally expensive.

**Results:** We developed ConsTrain—a gradient-based machine learning method for rich structural alignment scoring. We also implemented ConsAlign—a simultaneous RNA structural aligner composed of ConsTrain’s learned scoring parameters. To aim for better structural alignment quality, ConsAlign employs (1) transfer learning from well-defined scoring models and (2) the ensemble model between the ConsTrain model and a mature thermodynamic scoring model. Keeping comparable running time, ConsAlign demonstrated competitive alignment prediction quality among current RNA structural aligners.

**Availability and implementation:** Our code and our data are freely available at https://github.com/heartsh/consalign.

**Supplementary information:** Supplementary data are available at *Bioinformatics*

## 1 Introduction

Machine learning is applied to each significant bioinformatics subject, including structural bioinformatics. Also, machine learning has attracted computational scientists’ attention around folding single RNA sequences—or *mono-folding*. In mono-folding, modern parameter training methods optimize secondary structure scoring parameters based on both training RNA sequences and training RNA secondary structures (Do *et al*., 2006b; Zakov *et al*., 2011; Rivas *et al*., 2012; Singh *et al*., 2019; Sato *et al*., 2021). Folding RNA homolog sequences and aligning them at the same time— or *simultaneous RNA structural alignment* (Sankoff, 1985)— are an advanced solution, extending mono-folding. Simultaneous RNA structural alignment improves the folding quality of a focused RNA homolog sequence, incorporating other RNA homologs concerned. The machine learning methods of simultaneous RNA structural alignment have been developed less initiatively than those of mono-folding due to the complexity of the former (Dowell and Eddy, 2006; Do *et al*., 2008).

This study proposes *ConsTrain*—an *alignment score learning extension* of our sparse inside-outside algorithm (Tagashira and Asai, 2022). In Tagashira and Asai (2022), our sparse inside-outside algorithm scores possible simultaneous structural alignments by the scoring parameters of (1) Turner’s (nearest neighbor physics) model for RNA secondary structures (Turner and Mathews, 2010) and (2) the CONTRAlign model for RNA sequence alignments (Do *et al*., 2006b). By ConsTrain, we intend to improve the predictive performance of our sparse inside-outside algorithm implemented using these two independent models. ConsTrain is based on the *pairwise conditional log-linear model* (= pair-CLLM)—a flexible discriminative training framework (Do *et al*., 2006a). Scoring parameter training for simultaneous RNA structural alignment has been proposed using not the pair-CLLM but the pairwise stochastic context-free grammar (= the pair-SCFG)—a classical generative training framework (Dowell and Eddy, 2006). Regarding mono-folding accuracy, CLLMs are generally superior to SCFGs due to SCFGs’ model constraints (Do *et al*., 2006a).

To test the prediction performance of ConsTrain’s trained alignment parameters, we build *ConsAlign*—a simultaneous RNA structural aligner parameterized with them. ConsAlign conducts sparse dynamic programming according to the *progressive alignment scheme* (Feng and Doolittle, 1987). ConsAlign depends on its hyper-parameters to control simultaneous structural alignment quality but corrects them automatically by our original scheme without exploiting any validation data. As our first essential contribution, we introduce *transfer learning* (Singh *et al*., 2019) based on conventional trained models into ConsTrain. As our second essential contribution, we also introduce a *model ensemble* (Aghaeepour and Hoos, 2013; Singh *et al*., 2019) between our thermodynamic probability inference and ConsTrain’s trained probability inference into ConsAlign. We expect that ConsAlign will acquire higher precision and more robustness through our essential contributions.

## 2 Methods

### 2.1 Computing alignment scoring parameters’ gradient

We can optimize the scoring parameters of each CLLM (including our structural alignment pair-CLLM), given their cost functions (Do *et al*., 2006a; Andronescu *et al*., 2007). From both the empirical occurrence counts and the expected occurrence counts of pair-CLLM scoring parameters in observed pairwise structural alignments (Fig. 1), we can derive the *gradient* of a pair-CLLM cost function. We compute the approximated expected counts of scoring parameters appearing in training pairwise structural alignments by conducting our inside-outside algorithm on sparse pairwise structural alignment (Tagashira and Asai, 2022) iteratively.

**Fig. 1.**
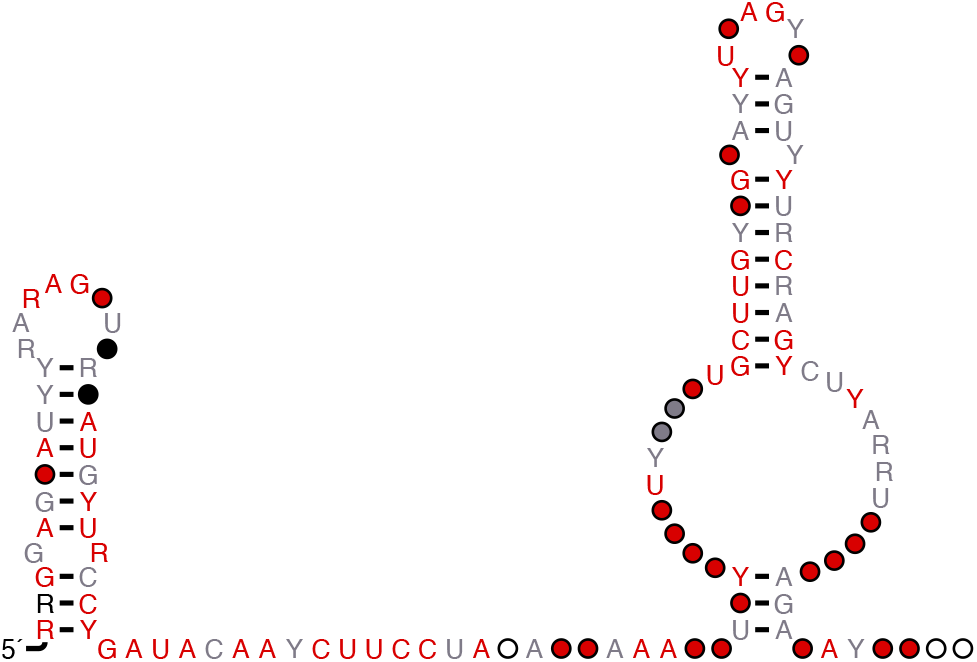
RNA structural alignment is depicted in R-scape (Rivas et al., 2017).

#### 2.1.1 Forming the cost of training pairwise structural alignments

We derive a *convex* pair-CLLM cost function fully parameterized with tunable alignment scoring parameters from training pairwise structural alignments. We score any pairwise structural alignment 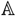 by using a simple inner product:

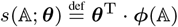

Here, ***θ*** is any scoring parameter vector in the real *F*-space. In addition, a function 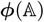 maps any pairwise structural alignment 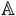 to the vector that counts the occurrence of each *f*-th scoring parameter *θ_f_* : (*θ_f_*) ≡ ***θ***, *f* ∈ {1,…, *F*} in 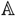. This study assumes that any structural alignment 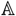 does not maintain any pseudoknot (Sato *et al*., 2011; Will *et al*., 2018) to simplify our proposed algorithms; we do not consider any base-pairing indels, i.e., we must match each base-pairing in one RNA secondary structure contained in 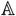 with some base-pairing in the other RNA secondary structure contained in 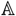, and vice versa, as implemented in many RNA structural aligners (Hofacker *et al*., 2004; Kiryu *et al*., 2007; Will *et al*., 2007; Do *et al*., 2008; Sato *et al*., 2012). The pairwise structural alignment scoring 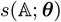 obeys an underlying pair-CLLM. As a result, each alignment scoring parameter *θ_f_* does not have to take the form of a log probability nor correspond only to a transition/emission (Do *et al*., 2006a).

Onto the training structural alignment 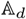 of each *d*-th RNA sequence pair, we obtain a convex pair-CLLM cost (Do *et al*., 2006a; Andronescu *et al*., 2007)

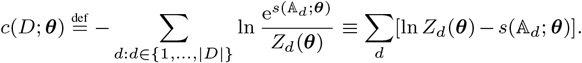

Here, any training alignment dataset *D* is 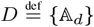. Furthermore, any alignment *partition function Z_d_*(***θ***) is 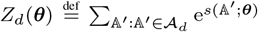. (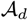 is the structural alignment *Boltzmann ensemble*—i.e., a set of all possible structural alignments—of each *d*-th sequence pair.) Our convex pair-CLLM cost *c*(*D*; ***θ***) holds its gradient

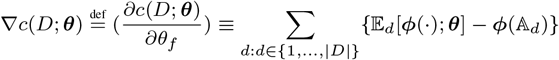

because any log function gradient 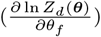 is

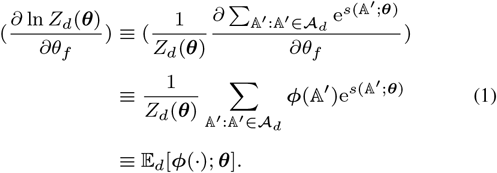

Here, any expected count vector 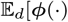 is

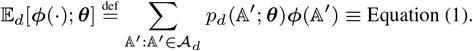

[Any alignment *Boltzmann (probability) distribution* 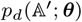 is 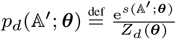.]

By using our sparse inside-outside algorithm, we can estimate the approximate amount of any expected count vector 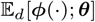 (Do *et al*., 2006a; Andronescu *et al*., 2007):

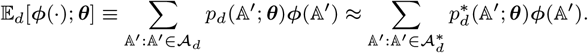

Here, 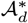 is a sparse version of any alignment Boltzmann ensemble 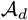.

In addition, any sparse Boltzmann distribution 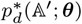 is as follows:

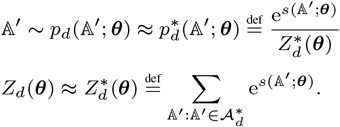

This approximated vector estimation requires the same running time *O*(*L*^2^) and memory usage *O*(*L*^2^) as our sparse inside-outside algorithm (Tagashira and Asai, 2022). Here, *L* is the longer length of each *d*-th sequence pair.

### 2.2 Optimizing structural alignment scoring parameters

Our parameter optimization goal is to find structural alignment scoring parameters that minimize their convex pair-CLLM cost. We can utilize general gradient-based optimization methods to complete our scoring parameter optimization. We aim to compute an optimized parameter vector

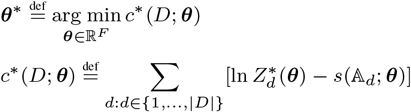

(where 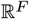 is the real *F*-space) using the gradient of an approximated pair-CLLM cost *c*\* (*D*; ***θ***)

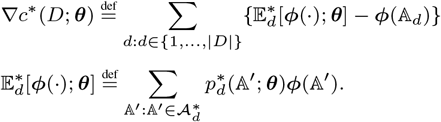

By Algorithm S1, we can conduct the above convex parameter optimization iteratively. Algorithm S1 is an instance of the Broyden-Fletcher-Goldfarb-Shanno (= BFGS) algorithm—a quasi-Newton optimization algorithm widely used (Fletcher, 1987).

Our approximated pair-CLLM cost *c*\*(*D*; ***θ***) does not retain any regularization term. In other words, an optimized parameter vector ***θ***\* is prone to be overfitted for each training alignment dataset *D*. Thus, we minimize an L2-regularized non-convex cost instead of our unregularized convex cost *c*\*(*D*; ***θ***):

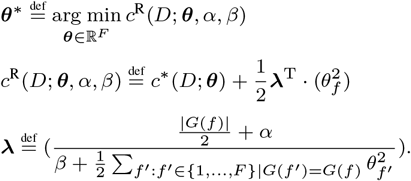

Here, *α, β* are the two gamma distribution parameters. Also, a function *G*(*f*) returns the group where each alignment scoring parameter *θ_f_* belongs. Our adaptive L2-regularization vector **λ** introduces a prior alignment parameter (probability) distribution:

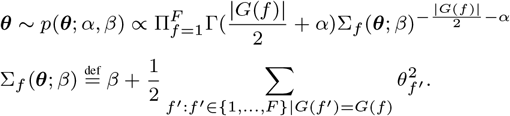

Here, the gamma function Γ(*r*) given any real number *r* is 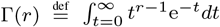. We implement the above non-convex parameter optimization iteratively by Algorithm S2 based on Algorithm S1. Algorithm S2 is an example of *majorization-minimization algorithm*s— iterative optimization algorithms that minimize some convex upper bound of any non-convex cost function (Langed *et al*., 2000). Algorithm S2 groups each alignment scoring parameter *θ_f_* into a cluster and penalizes each scoring parameter cluster by a shared regularization constant (Foo *et al*, 2009). The advantage of introducing our prior parameter distribution *p*(***θ***; *α, β*) is that we do not have to predetermine the value of our L2-regularization constant vector **λ** (e.g., by a grid search upon a validation structural alignment dataset) since we (in addition to our alignment scoring parameters **θ** to be optimized) optimize **λ** in the same procedure (Foo *et al*, 2009). We implemented Algorithm S2 as the ConsTrain version 0.1.1 (https://github.com/heartsh/consprob-trained).

### 2.3 Configuring structural alignment scoring parameters

Borrowing knowledge from conventional scoring parameter optimizations, we specify the concrete settings of our alignment scoring parameters. We

specify all the groups (or types) of our alignment scoring parameters, as described in Table S1. The *CONTRAfold model* is a CLLM that (from the training structure of each single RNA sequence) learns secondary structure scoring parameters based on RNA loop structures (Do *et al*., 2006a). The *CONTRAlign model* is a pair-conditional random field (Lafferty *et al*., 2001; Sato and Sakakibara, 2005) that trains sequence alignment scoring parameters from each sequence-aligned RNA pair (Do *et al*., 2006b). Our pair-CLLM inherits scoring parameter formats from both the CONTRAfold and the CONTRAlign models.

### 2.4 Our proposed structural aligner to maximize accuracy

#### 2.4.1 Quadratic pairwise alignment guaranteeing its optimality

In the γ-centroid estimator principle (Ding *et al*., 2005; Hamada *et al*., 2009a,b,c, 2011), we define a pairwise structural alignment recursion like LocARNA (Will *et al*., 2007) and RAF (Do *et al*., 2008), as in Fig. 2:

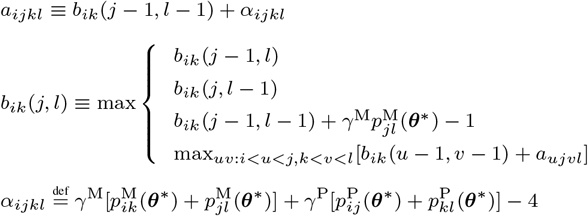

starting from the initial condition *b_ik_*(*i,k*) ← 0. Here, 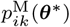 is the sparse posterior probability that any nucleotide (position) pair (*i, k*) matches in pairwise structural alignment given our trained scoring parameters ***θ***\*; 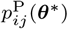 is the average sparse probabilistic consistency that every two nucleotides *i, j* base-pair in pairwise structural alignment given ***θ***\*. The new ConsProb (Tagashira and Asai, 2022) parameterized with our trained scoring parameters ***θ***\* can compute trained sparse probabilities 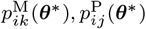. Also, *γ*^M^, *γ*^P^ are any two real numbers greater than 1. Hyper-parameters *γ*^M^ and *γ*^P^ control the predictive significance of nucleotide matching and nucleotide base-pairing, respectively.

**Fig. 2.**
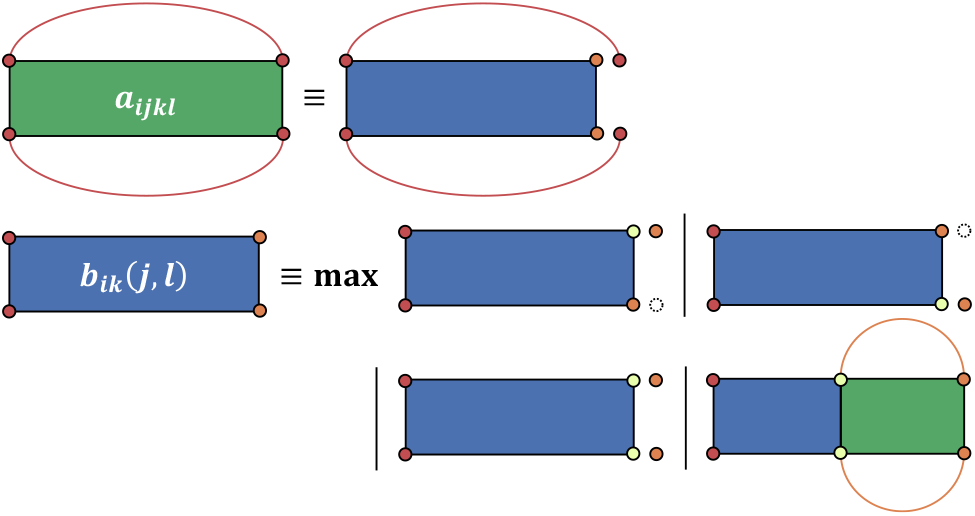
Recursive diagram of our pairwise structural alignment.

We need to conduct our pairwise predictive recursion with *O*(*N*^3^*M*^3^) running time and *O*(*N*^2^*M*^2^) memory usage in a naive sense. (N, M are the lengths of every two RNA sequences.) However, we can perform our pairwise predictive recursion only with both *O*(*L*^2^) running time and *O*(*L*) memory usage if hyper-parameters *γ*^M^, *γ*^P^ are sufficiently small. [*L* is *L* ≝ max(*N, M*).] We can realize this computational complexity reduction since our pairwise predictive recursion accepts only nucleotide matches (*i, k*) satisfying the condition

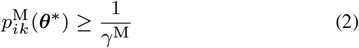

and nucleotide base-pairings (*i, j*), (*k, l*) satisfying the conditions

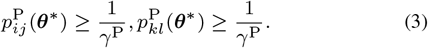

LocARNA and RAF impose Equations (2) and (3) upon possible pairwise structural alignments to enable both *O*(*L*^2^) running time and *O*(*L*) memory usage (Do *et al*., 2008). Our pairwise predictive recursion assures that its obtained structural alignment is optimum (i.e., retaining the best score of all possible structural alignments) even if we impose Equations (2) and (3) upon possible pairwise structural alignments.

#### 2.4.2 Extending pairwise alignment into multiple-alignment

Our structural multiple-alignment is based on the progressive alignment scheme (Feng and Doolittle, 1987) like conventional RNA structural aligners (Will *et al*., 2007; Do *et al*., 2008; Sato *et al*., 2012; Will *et al*., 2015; Li *et al*., 2021). When we build a *UPGMA guide tree* (Sneath and Sokal, 1962) from RNA homolog sequences to be aligned, we measure the efficient similarity between every two RNA homologs

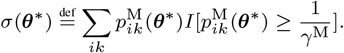

Here, *I*[*c*] returns 1 if some condition *c* is true; otherwise, *I*[*c*] returns 0. Given a hyper-parameter *γ*^M^, our proposed metric *σ*(***θ***\*) estimates the stochastic similarity between every two RNA homologs without aligning them. Our proposed metric *σ*(***θ***\*) considers only nucleotide matches (*i, k*) satisfying the condition 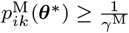 to be eligible.

We extend our pairwise predictive recursion into a generalized multiple predictive recursion, replacing pairwise nucleotide matching probabilities 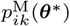 with alignment column matching probabilities

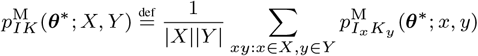

and replacing alignment-based average base-pairing probabilities 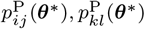 with alignment column base-pairing probabilities

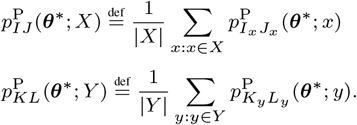

Here, *X, Y* are every two aligned sets of RNA homolog sequences to be aligned. Moreover, *I, J, K, L* are every four alignment columns, and *I_x_* is any RNA homolog *x*’s nucleotide stored in *I*. 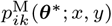 is the explicit form of each nucleotide matching probability 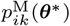 regarding every two RNA homologs *x, y*; and 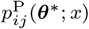 is the explicit form of each average base-pairing probability 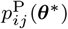 regarding *x*.

#### 2.4.3 Auto-correction of structural alignment hyper-parameters

The disadvantage of *γ*-centroid estimators is that they rely on hyperparameters (e.g., hyper-parameters *γ*^M^, *γ*^P^ in our structural alignment). In practice, we have to determine hyper-parameters before performing *γ*-centroid estimators. This study proposes a reasonable way to determine hyper-parameters in performing *γ*-centroid simultaneous structural alignments. The *sum-of-pairs score* (= SPS) of any predicted structural alignment 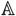 against any possible multiple alignment 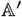 is

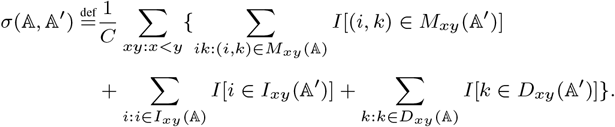

Here, an SPS denominator *C* is

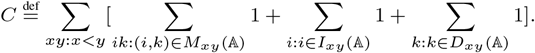

In addition, 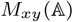 is a set of all aligned nucleotide pairs between every two RNA homologs *x,y* in any structural alignment 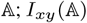 is a set of all inserted nucleotides between *x, y* in 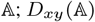 is a set of all deleted nucleotides between *x, y* in 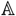.

As a result, we can measure the approximated expected SPS of any predicted structural alignment 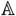 against all possible multiple alignments

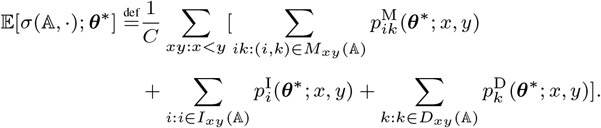

Here, 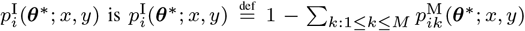 and 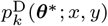 is 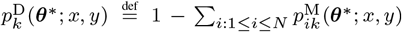. Finally, we can optimize hyper-parameters *γ*^M^, *γ*^P^ as well as a predicted structural alignment by enumerating possible values of *γ*^M^, *γ*^P^ as in Algorithm S3. Our alignment hyper-parameter auto-correction is similar to hyper-parameter auto-correction used in mono-folding—called pseudo-expected accuracy optimization (Hamada *et al*., 2010; Sato and Kato, 2022)—because both these methods optimize *γ*-centroid hyper-parameters based on posterior probabilities.

#### 2.4.4 *Ali-folding* post-process of predicted structural alignment

Folding RNA alignment—or ali-folding—is one of the most potent ways to find *conserved base pairings* in RNA alignment as well as simultaneous RNA structural alignment (Bernhart *et al*., 2008; Seemann *et al*., 2008; Hamada *et al*., 2011; Tagashira and Asai, 2022). As LocARNA adopts RNAalifold (Bernhart *et al*., 2008) as the ali-folding post-processor of LocARNA’s predicted structural alignment (Will *et al*., 2007), we adopt ConsAlifold (Tagashira and Asai, 2022) parameterized with our trained alignment parameters as the ali-folding post-processor of our predicted structural alignment.

#### 2.4.5 Ensemble of different alignment scoring models

Unifying different stochastic models for a prediction—referred to as a model ensemble—has been utilized in mono-folding to achieve more accurate structure prediction (Aghaeepour and Hoos, 2013; Singh *et al*., 2019). In our structural alignment prediction, we combine our sparse inside-outside algorithm of Tagashira and Asai (2022) and that of this study to create an alignment scoring model ensemble. More specifically, we use average column matching probabilities and average column base-pairing probabilities

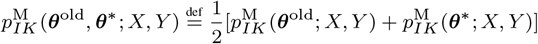

and average column base-pairing probabilities

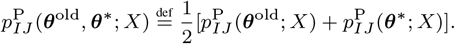

Here, ***θ***^old^ is the alignment scoring parameters used in Tagashira and Asai (2022). Our old scoring parameters ***θ***^old^ are from both Turner’s and the CONTRAlign models, while our trained scoring parameters ***θ***\* are from the CONTRAfold and the CONTRAlign models. We implemented our RNA structural aligner explained above as the ConsAlign version 0.1.1 (https://github.com/heartsh/consalign).

### 2.5 Benchmarking data

We used the Rfam version 14.4 (Kalvari *et al*., 2018) for our alignment parameter training and software benchmarking test. From Rfam, we collected 3129 structurally aligned families as the dataset “origin.” The seed reference structural alignment of each collected RNA family retained at most 500 columns and 20 sequences. We randomly split the dataset “origin” into both the equal datasets “train origin” and “test multiple.” From the dataset “train origin,” we collected each RNA family whose seed reference consensus structure is not any prediction product [e.g., neither by Pfold (Knudsen and Hein, 2003) nor by RNAalifold (Bernhart *et al*., 2008)] as the dataset “train pure.” We random-sampled 10 RNA sequences from each RNA family holding at least 10 RNA sequences in the dataset “train pure.” Then, we retrieved the all-to-all pairwise structural alignments of these 10 RNA sequences, finally obtaining the dataset “train pairwise.” This random sequence sampling prevents our alignment parameter learning from overfitting due to the overwhelming number of the RNA homologs belonging to the same large RNA family (Do *et al*., 2006a, 2008).

As a result, (1) the dataset “train pairwise” is composed of 2250 pairwise structural alignments, and (2) the dataset “test multiple” is composed of 1565 multiple structural alignments. We used the dataset “train pairwise” in order to train ConsTrain. We used the dataset “test multiple” to assess the prediction performances of each benchmarked prediction method. Any entry in the dataset “train pairwise” and any entry in the dataset “test multiple” are not derived from the same family to prevent structural homology from leaking between training data and test data (Rivas *et al*., 2012; Sato *et al*., 2021).

As another test data source, we adopted reference RNA structural alignments registered in RNAStralign (Tan *et al*., 2017). We split each complete structural alignment comprising all RNA sequences into the subalignment comprising every 20 or fewer sequences. At last, we obtained the dataset “RNAStralign processed,” composed of 917 multiple structural alignments. From the BRAliBase version 2 (Wilm *et al*., 2006), we also downloaded dataset 1, containing reference RNA sequence alignments, as a test dataset.

### 2.6 Conventional structural aligners for experiments

Using default command line arguments, we compared (1) the RAF version 1.00, (2) the LocARNA version 2.0.0, (3) the SPARSE version 2.0.0 (Will *et al*., 2015), (4) the DAFS version 0.0.3 (Sato *et al*., 2012), and (5) the LinearTurboFold version 1 (Li *et al*., 2021) to ConsAlign. Each conventional aligner’s characteristics are described in Section S4.

### 2.7 Alignment prediction accuracy metrics

Across all test structural alignments, we find the numbers of true positives, true negatives, false positives, and false negatives *TP, TN, FP, FN* regarding structure quality while benchmarking each tool. We compute an F1 score and a Matthews correlation coefficient (= MCC) in each test alignment dataset:

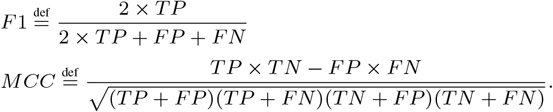

To measure the matching quality of each predicted structural alignment, we obtain both the SPS and the structure conservation index (= SCI) of each predicted structural alignment. Each SCI quantifies the structural conservation of a corresponding RNA sequence alignment from a thermodynamic viewpoint (Washietl *et al*., 2005) and, therefore, diagnoses the matching quality of each predicted RNA alignment considering potential structure quality.

### 2.8 Implementations and benchmark environments

Employing multi-threading to provide a more comfortable user experience, we implemented ConsTrain and ConsAlign in Rust. ConsTrain used the two gamma distribution parameters *α* ← 0, *β* ← 1 proposed by Foo *et al* (2009). In ConsTrain, we implemented two different initialization methods for structural alignment scoring parameters to be trained. (1) In our first initialization method, corresponding scoring parameter values were transferred from the CONTRAfold and the CONTRAlign models. (2) In the other initialization method, structural alignment scoring parameters were initialized by random values derived from the normal distribution with both the mean 0 and the variance 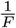. ConsAlign used the four lower/upper bounds *i^min^* ← 0, *i*^max^ ← 7, *j*^min^ ← 0, *j*^max^ ← 3 found in Algorithm S3. Also, ConsAlifold performed in ConsAlign took the γ-centroid hyper-parameter 2^0^ + 1 = 2 because this γ-centroid hyperparameter value showed the highest structure prediction accuracy among the possible γ-centroid hyper-parameter values 2^−4^ + 1, …, 2^10^ + 1 in Tagashira and Asai (2022). For our prediction software benchmarks, we used an “Intel Xeon” CPU computer with 240 GB of RAM, 64 threads, and a clock rate of 2.30 GHz. (We refer to this computer as the environment “Xeon.”)

## 3 Results

### 3.1 Benchmarking ConsAlign to conventional methods

ConsAlign consistently performed better structure prediction accuracy than the other RNA structural aligners, whereas LinearTurboFold (Li *et al*., 2021) had the lowest rank regarding secondary structure prediction accuracy (Fig. 3a). ConsAlign showed comparable SPSs among state-of-the-art RNA structural aligners, while SPARSE (Will *et al*., 2015) indicated the poorest SPSs (Fig. 3b). Regarding SCIs, we did not observe any significant difference among all compared structural aligners except LinearTurboFold (Fig. 3b). ConsAlign took the median prediction running time among available RNA structural aligners (Table 1).

**Fig. 3.**
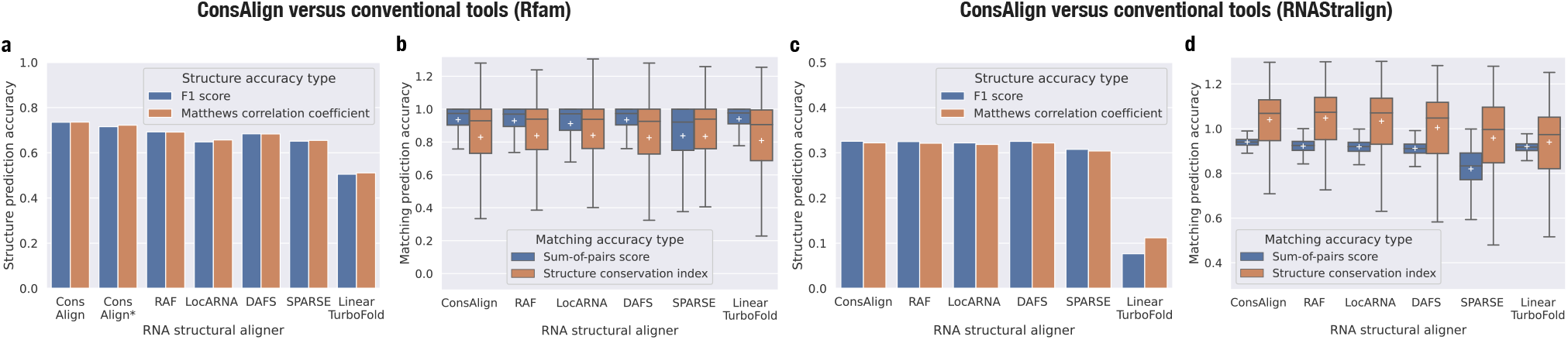
Alignment prediction accuracy comparison among ConsAlign and conventional structural aligners. (a, c) Methods were compared in terms of F1 scores and MCCs. (b, d) Each method’s distributions of SPSs and SCIs are visualized in box plots. Outliers were omitted from box plots. Each white cross indicates a corresponding method’s average. The datasets (a, b) “test multiple” and (c, d) “RNAStralign processed” were used. ConsAlign was based on our proposed ensemble model and our proposed transfer learning. (ConsAlign*) ConsAlign disabled its RNAalifold subroutine from producing alignment column base-pairing probabilities (Bernhart et al., 2008) to check ConsAlign’s pure prediction performance.

**Table 1.** Prediction type and running time of benchmarked structural aligners.

| Structural aligner | Structural alignment type | Total running time |
| --- | --- | --- |
| ConsAlign | Simultaneous | 442 s |
| RAF |  | 1697 s |
| LocARNA | : | 70 s |
| SPARSE |  | 618 s |
| DAFS | Simultaneous | 84 s |
| LinearTurboFold | Iterative | 1362 s |

The center column indicates each aligner’s prediction type. The right column shows the total running time of each benchmarked structural aligner on the dataset “test multiple.” Utilizing `multiprocessing.Pool` (https://docs.python.org/3/library/multiprocessing.html) to relieve the large volume of the dataset “test multiple,” each benchmarked structural aligner other than ConsAlign was run on the environment “Xeon” by mapping 64 processes. ConsAlign was based on our proposed ensemble model and our proposed transfer learning. Moreover, ConsAlign was conducted by providing each of 16 processes with four different threads.

When we switched our test dataset from the dataset “test multiple” to the dataset “RNAStralign processed,” ConsAlign competed with most RNA structural aligners regarding predicted structures’ quality, separating LinearTurboFold from the others (Fig. 3c). ConsAlign was superior to conventional RNA structural aligners from the viewpoint of SPS, and SPARSE suffered lower SPSs compared to the others (Fig. 3d). The SCIs of ConsAlign were comparable to those of RAF, LocARNA, and DAFS (Fig. 3d). Surprisingly, LinearTurboFold was more disadvantageous to SCI than the others, albeit performing moderate SPSs (Fig. 3d).

In BRAliBase’s dataset 1, ConsAlign and the existing structural aligners other than SPARSE were almost equally precise regarding SPSs (Fig. 4a–b). However, SPARSE’s predicted structural alignments were graded highly together with RAF and LocARNA in terms of SCIs (Fig. 4a,c).

**Fig. 4.**
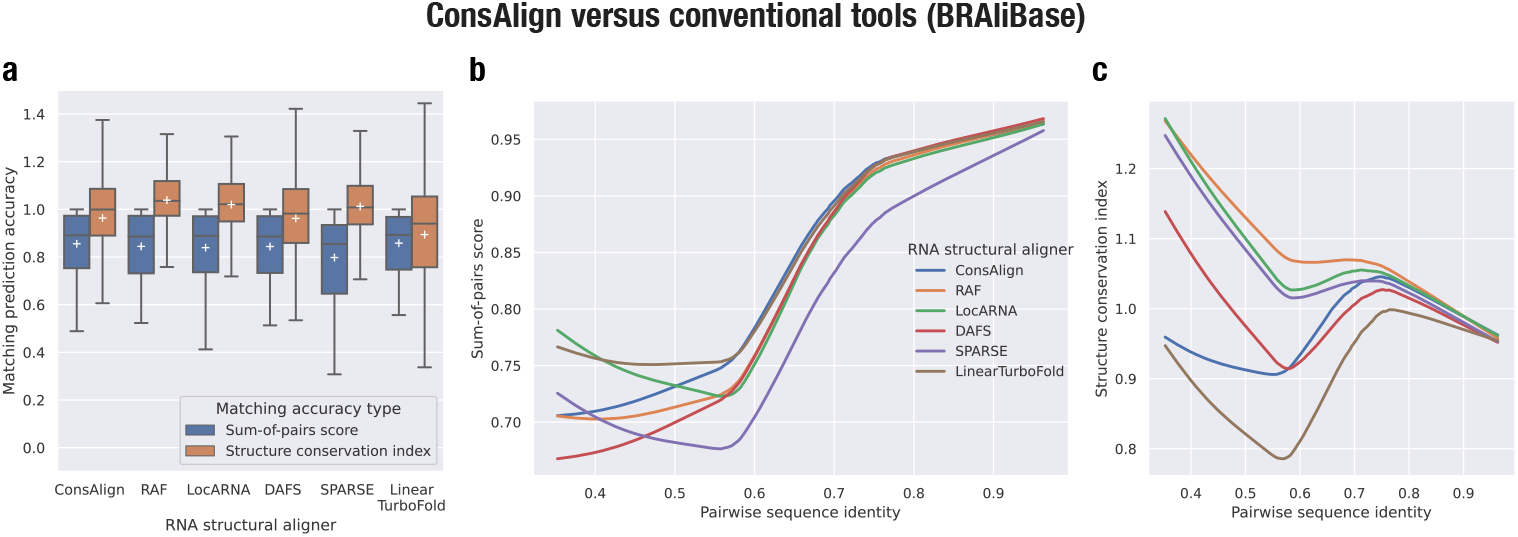
Predicted alignments’ quality among modern structural aligners. (a) Each method’s SPSs and SCIs are visualized in box plots. Each method’s (b) SPSs and (c) SCIs are expressed in lowess curves (Cleveland, 1981) to the pairwise sequence identity of reference alignments. BRAliBase’s dataset 1 was used.

**Fig. 5.**
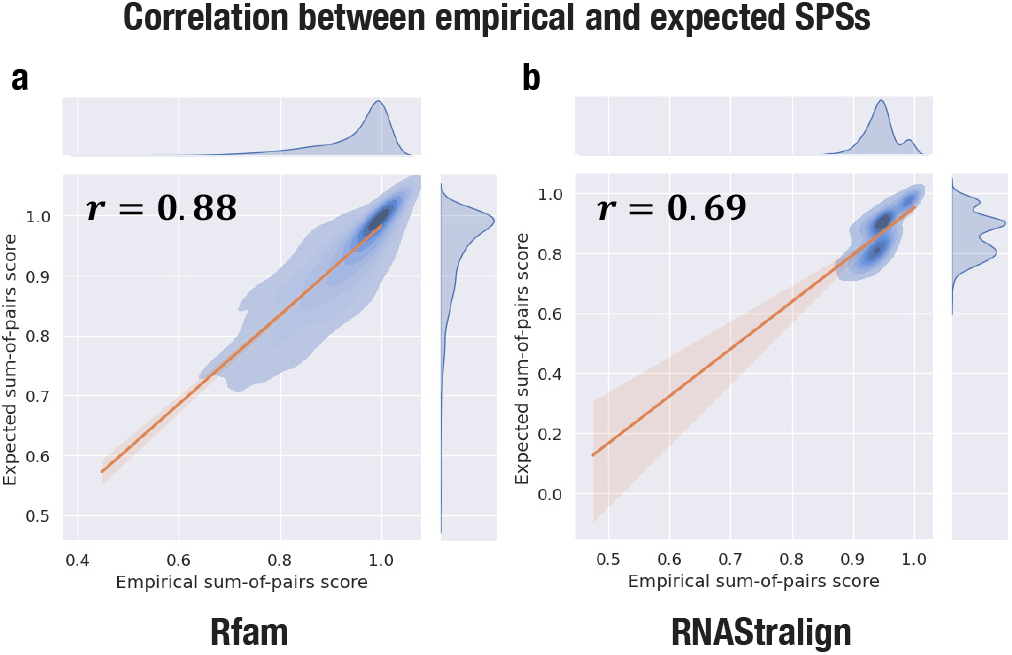
Contrast between ConsAlign’s empirical and expected SPSs in the datasets (a) “test multiple” and (b) “RNAStralign processed.” (*r*) Each Pearson correlation coefficientwas noted. ConsAlign was based on our proposed ensemble model and our proposed transfer learning.

**Fig. 6.**
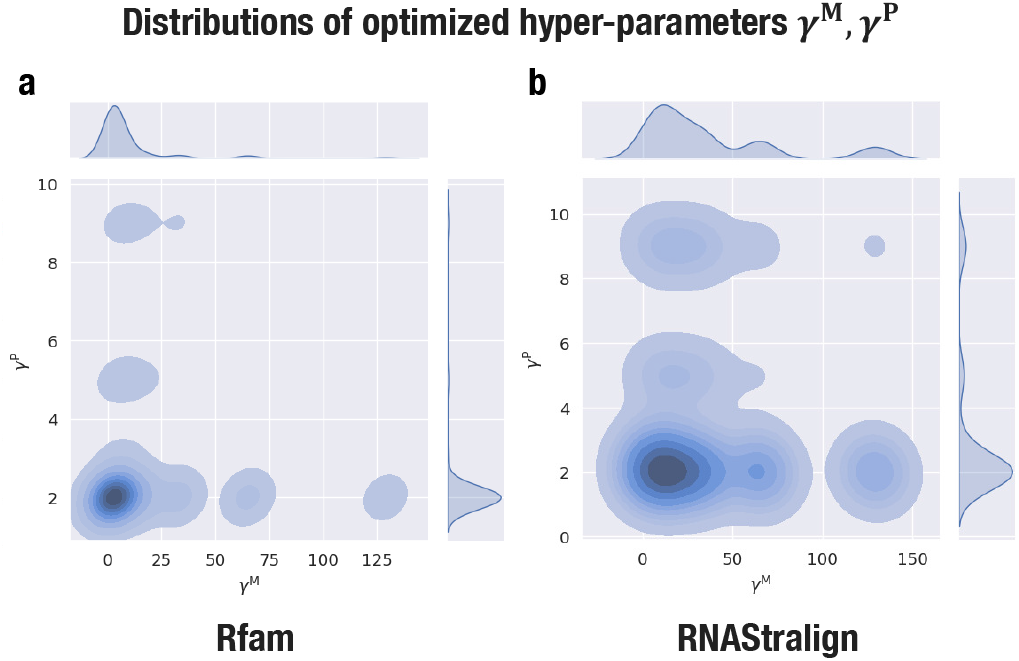
Distributions of ConsAlign’s optimized hyper-parameters *γ*^M^, *γ*^P^ in the datasets (a) “test multiple” and (b) “RNAStralign processed.” ConsAlign was based on our proposed ensemble model and our proposed transfer learning.

### 3.2 Verifying our proposed training and model ensemble

ConsTrain took only about 30 epochs when using our proposed transfer learning (Fig. S1). On the other hand, ConsTrain consumed around 60 epochs when we initialized our alignment scoring parameters to random values (Fig. S1). ConsTrain continuously reduced our L2-regularized non-convex cost in both parameter initialization methods (Fig. S1). In each test dataset, ConsAlign’s expected SPSs highly correlated with its observed SPSs (Figs. S2 and 5), and ConsAlign’s tuned hyper-parameters were principally distributed around lower bounds (Figs. S2 and 6).

In the dataset “test multiple,” ConsTrain with our proposed transfer learning demonstrated the best structure prediction accuracy among available parameter training schemes (Fig. 7a). Demonstrated parameter training schemes showed the same level of predicted matches’ quality (Fig. 7b). Our proposed ensemble model drew advantages from the old ConsProb, which outperformed the new ConsProb, in structural accuracy (Fig. 8a). In contrast, performed alignment scoring models were nearly equivalent for SPS and SCI (Fig. 8b).

**Fig. 7.**
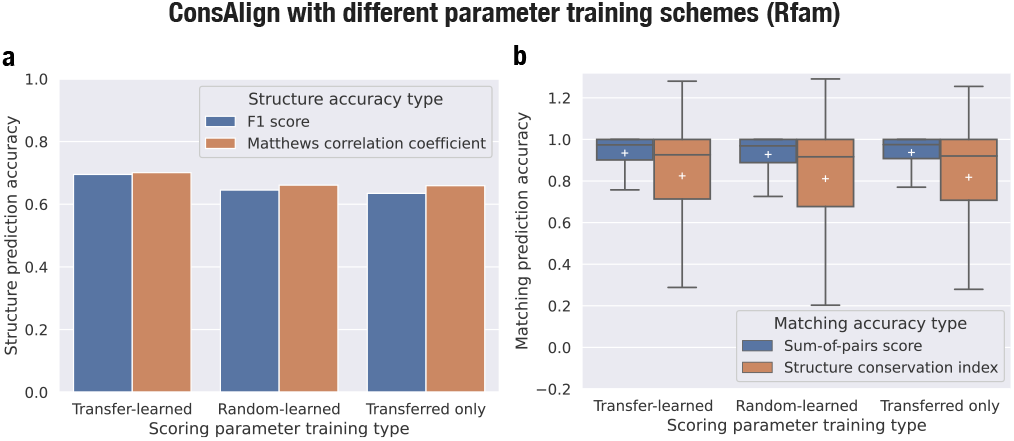
Alignment prediction accuracy comparison among the different parameter training schemes of ConsTrain. (Transfer-learned) Structural alignment scoring parameters were transfer-learned. (Random-learned) Random values initialized structural alignment scoring parameters before training. (Transferred only) Structural alignment scoring parameters were transferred but then were not trained. This figure’s configuration is the same as Fig. 3. The dataset “test multiple” was used. ConsAlign was based on the new ConsProb (Tagashira and Asai, 2022) alone and disabled an RNAalifold subroutine.

**Fig. 8.**
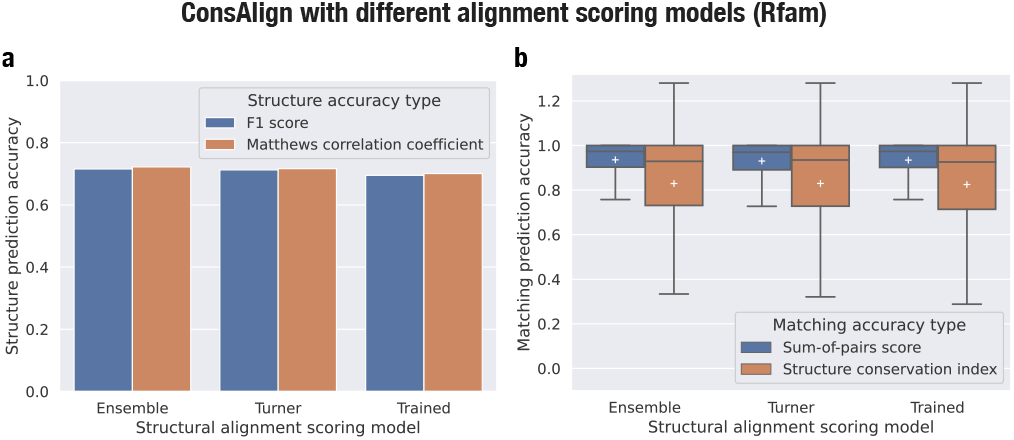
Alignment prediction accuracy comparison among the different alignment scoring models of ConsAlign. (Ensemble) Our proposed ensemble model was used. (Turner) Only the old ConsProb was used as a posterior probability inference method. (Trained) Only the new ConsProb was used as a posterior probability inference method. This figure’s configuration is the same as Fig. 3. The dataset “test multiple” was used. ConsAlign was based on our proposed transfer learning and disabled an RNAalifold subroutine.

Shifting to the dataset “RNAStralign processed,” ConsAlign was not enhanced at any alignment quality criterion when we focused on our transfer learning against our scoring parameters that were only transferred (Fig. S3). Regarding both structural and matching quality, our proposed ensemble model capitalized on the new ConsProb, which surpassed the old ConsProb (Fig. S4). From BRAliBase’s dataset 1, predicted matches’ quality was raised in ConsAlign when we compared our transfer learning with the others (Fig. 9a–b, Fig. S5a). Our proposed ensemble model used the new ConsProb and the old ConsProb to perform the best SPSs and SCIs (Fig. 9c–d, Fig. S5b).

**Fig. 9.**
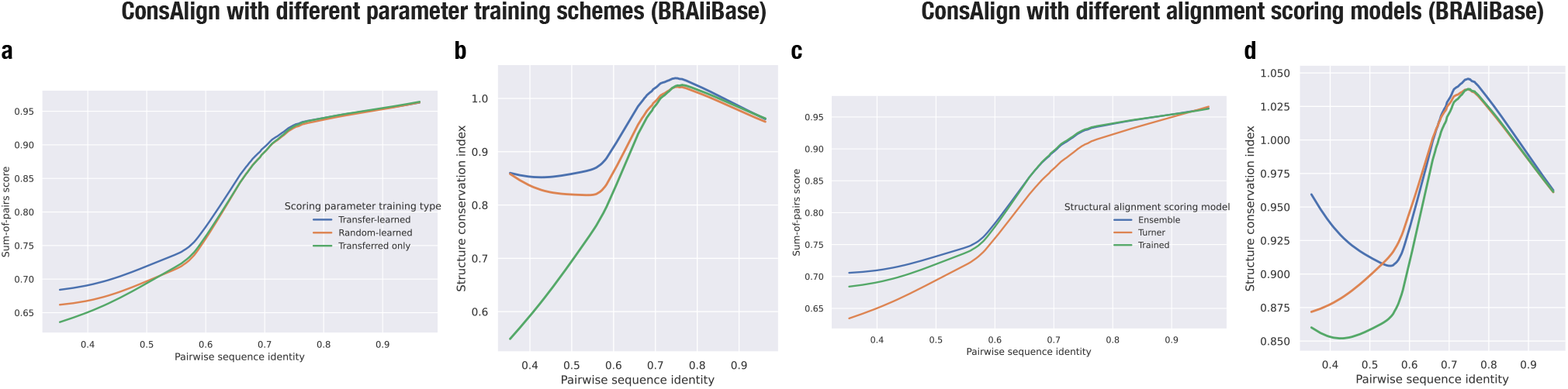
Alignment prediction accuracy comparison among (a, b) the different parameter training schemes and (c, d) the different alignment scoring models of ConsAlign. This figure’s configuration is the same as Figs. 7 and 8. BRAliBase’s dataset 1 was used.

## 4 Conclusion and discussion

We invented ConsTrain—an efficient gradient-based learning method of simultaneous alignment scoring parameters—and applied it to ConsAlign—a model ensemble-based structural aligner. ConsAlign attained competitive alignment quality in our alignment tool benchmarks. In our alignment software comparison, LinearTurboFold and SPARSE were the weakest in structural prediction accuracy and SPSs, respectively. LinearTurboFold is an iterative method different from the other benchmarked simultaneous methods. Therefore, we can interpret that simultaneous solutions enhanced obtained structures’ quality by the more direct transmission of structural conservation among homologs than iterative solutions. SPARSE realizes its efficient computation by employing only structure-based constraints, whereas the other simultaneous structural aligners—ConsAlign, LocARNA, RAF, and DAFS—employ matching-based constraints in addition to structure-based constraints. We can explain that leveraging both matching-based structure-based constraints is more effective regarding obtained nucleotide matches’ quality than only structure-based constraints.

ConsTrain with our transfer learning substantiated double parameter training efficiency and better structural alignment quality than our random parameter initialization. ConsTrain’s iterative parameter update is not convex due to our attached training hyper-parameters. Thus, we can conclude that providing well-defined scoring parameter values guided ConsTrain to better machine learning solutions. Our proposed ensemble model made the flexible use of both the new ConsProb and the old ConsProb to achieve ConsAlign’s robust alignment quality. In other words, both thermodynamics-based and entirely statistical models compensated each other for predictive weak points.

Recently, deep learning-based methods for mono-folding—such as SPOT-RNA (Singh *et al*., 2019), MXfold (Sato *et al*., 2021), and Ufold (Fu *et al*., 2022)—emerged, replacing non-deep learning methods for monofolding—such as CONTRAfold (Do *et al*., 2006b) and TORNADO (Rivas *et al*., 2012). Expanding ConsTrain into deep learning for simultaneous RNA structural alignments has the potential to improve their quality by in-depth features unable to capture in any CLLMs. In another scenario, we have a chance to generalize the standard RNA base alphabet (`A, C, G`, and `U`) used in ConsTrain by including modified bases—such as `I` and Ψ (Harcourt *et al*., 2017). However, we need well-defined training datasets containing modified bases in this scenario because the base composition of training datasets directly affects ConsTrain’s trained scoring parameters via gradients. Constructing reliable large structural databases for modified bases can assist machine learning methods involving RNA secondary structures in data-driven avenues.

## Acknowledgments

We would like to thank Dr. Kiyoshi Asai, Dr. Martin Frith, and their lab members for a year of discussion. We incredibly appreciate valuable comments from Dr. Kengo Sato and Dr. Akito Taneda regarding points to be improved. We performed most of the computations in this study on the NIG supercomputer at the ROIS National Institute of Genetics, Japan.

## Data Availability

Our underlying data are available at our shown GitHub repositories.

