## Supplementary material for "ConsAlign: simultaneous RNA structural aligner based on rich transfer learning and thermodynamic ensemble model of alignment scoring"

### S1 Table

Table S1. Specified types of our alignment scoring parameters.

| Alignment scoring parameter group | Group size | Source |
|---|---|---|
| Hairpin loop length | 31 | CONTRAfold |
| Bulge loop length | 30 | |
| Interior loop length | 29 | |
| Interior loop length symmetric | 15 | |
| Interior loop length asymmetric | 28 | |
| Stacking | 36 | |
| Terminal mismatch | 96 | |
| Left dangling | 24 | |
| Right dangling | 24 | |
| | | ⋮ |
| Helix end | 6 | |
| Base-pairing | 6 | |
| Interior loop length explicit | 16 | |
| Bulge loop length $0 \times 1$ | 4 | |
| Interior loop length $1 \times 1$ | 16 | |
| Multi-loop base | 1 | |
| Multi-loop base-pairing | 1 | |
| Multi-loop accessible unpairing | 1 | |
| External-loop accessible base-pairing | 1 | |
| External-loop accessible unpairing | 1 | CONTRAfold |
| Match-to-match transition | 1 | CONTRAlign |
| Match-to-insert transition/insert-to-match transition | 1 | |
| Insert switch transition | 1 | |
| | | ⋮ |
| Initial match transition | 1 | |
| Initial insert transition | 1 | |
| Insert extension transition | 1 | |
| Match emission | 16 | |
| Insert emission | 4 | CONTRAlign |
| Total (from 27 groups) | 392 | |

Our structural alignment pair-CLLM combines the scoring parameter forms of the CONTRAfold model and the CONTRAlign model. Our pair-CLLM assumes the standard RNA nucleotide alphabet `A`, `C`, `G`, `U` and the canonical RNA base-pairing alphabet `AU`, `CG`, `GU`, `UA`, `GC`, `UG`.

### S2 Algorithms

**Algorithm S1** BFGS algorithm that minimizes an approximated convex pair-CLLM cost $c^*(D; \boldsymbol{\theta})$.

**function** bfgsAlgo(a training structural alignment dataset $D$)
  Initialize structural alignment scoring parameters $\boldsymbol{\theta}^*$
  **for** $d \in \{1, \dots, |D|\}$ **do**
    Get parameter counts $\boldsymbol{\phi}(\mathbb{A}_d)$ by parsing each training structural alignment $\mathbb{A}_d$
  **while** $\boldsymbol{\theta}^*$ does not converge **do**
    **for** $d \in \{1, \dots, |D|\}$ **do**
      Compute expected counts $\mathbb{E}^*_d[\boldsymbol{\phi}(\cdot); \boldsymbol{\theta}^*]$ using our inside-outside algorithm
    $\nabla c^*(D; \boldsymbol{\theta}^*) \leftarrow \sum_{d:d \in \{1,\dots,|D|\}} \{\mathbb{E}^*_d[\boldsymbol{\phi}(\cdot); \boldsymbol{\theta}^*] - \boldsymbol{\phi}(\mathbb{A}_d)\}$
    /* Apply the parameter update scheme of the BFGS algorithm */
    $\boldsymbol{\theta}^* \leftarrow \text{argmin}^{\text{BFGS}}[c^*(D; \boldsymbol{\theta}^*) | \nabla c^*(D; \boldsymbol{\theta}^*)]$
  **return** $\boldsymbol{\theta}^*$

**Algorithm S2** Grouped majorization-minimization algorithm that minimizes an L2-regularized non-convex cost $c^{\text{R}}(D; \boldsymbol{\theta}, \alpha, \beta)$.

**function** groupedMmAlgo(a training structural alignment dataset $D$, the two gamma distribution parameters $\alpha, \beta$)
  Initialize structural alignment scoring parameters $\boldsymbol{\theta}^*$
  **while** $\boldsymbol{\theta}^*$ does not converge **do**
    $\nabla c^*(D; \boldsymbol{\theta}^*) \leftarrow \sum_{d:d \in \{1,\dots,|D|\}} \{\mathbb{E}^*_d[\boldsymbol{\phi}(\cdot); \boldsymbol{\theta}^*] - \boldsymbol{\phi}(\mathbb{A}_d)\}$
    $\nabla c^{\text{R}}(D; \boldsymbol{\theta}^*, \alpha, \beta) \leftarrow \nabla c^*(D; \boldsymbol{\theta}^*) + \boldsymbol{\lambda}^{\text{T}} \cdot \boldsymbol{\theta}^*$
    $\boldsymbol{\theta}^* \leftarrow \text{argmin}^{\text{BFGS}}[c^{\text{R}}(D; \boldsymbol{\theta}^*, \alpha, \beta) | \nabla c^{\text{R}}(D; \boldsymbol{\theta}^*, \alpha, \beta)]$
    /* $(\theta^*_f) \stackrel{\text{def}}{=} \boldsymbol{\theta}^*$ */
    $\boldsymbol{\lambda} \leftarrow (\frac{\frac{|G(f)|}{2} + \alpha}{\beta + \frac{1}{2} \sum_{f':f' \in \{1,\dots,F\} | G(f') = G(f)} (\theta^*_{f'})^2})$
  **return** $\boldsymbol{\theta}^*$

**Algorithm S3** Algorithm that optimizes hyper-parameters $\gamma^{\text{M}}, \gamma^{\text{P}}$ together with a predicted structural alignment.

**function** alignerWrapper(RNA homolog sequences to be aligned, structural alignment scoring parameters $\boldsymbol{\theta}^*$)
  **for** $i \in \{i^{\min}, \dots, i^{\max}\}, j \in \{j^{\min}, \dots, j^{\max}\}$ **do**
    $\gamma^{\text{M}} \leftarrow 2^i + 1, \gamma^{\text{P}} \leftarrow 2^j + 1$
    Predict an alignment $\mathbb{A}$ using the progressive alignment scheme, given $\gamma^{\text{M}}, \gamma^{\text{P}}$
    Compute the expected sum-of-pairs score $\mathbb{E}[\sigma(\mathbb{A}, \cdot); \boldsymbol{\theta}^*]$ of $\mathbb{A}$
    $\sigma^{\text{new}} \leftarrow \mathbb{E}[\sigma(\mathbb{A}, \cdot); \boldsymbol{\theta}^*]$
    **if** $\sigma^{\text{new}} > \sigma^{\text{curr}}$ or $\sigma^{\text{curr}}$ is unset **then**
      $\mathbb{A}^* \leftarrow \mathbb{A}, \gamma^{\text{M},*} \leftarrow \gamma^{\text{M}}, \gamma^{\text{P},*} \leftarrow \gamma^{\text{P}}, \sigma^{\text{curr}} \leftarrow \sigma^{\text{new}}$
  **return** $(\mathbb{A}^*, \gamma^{\text{M},*}, \gamma^{\text{P},*})$

### S3 Figures

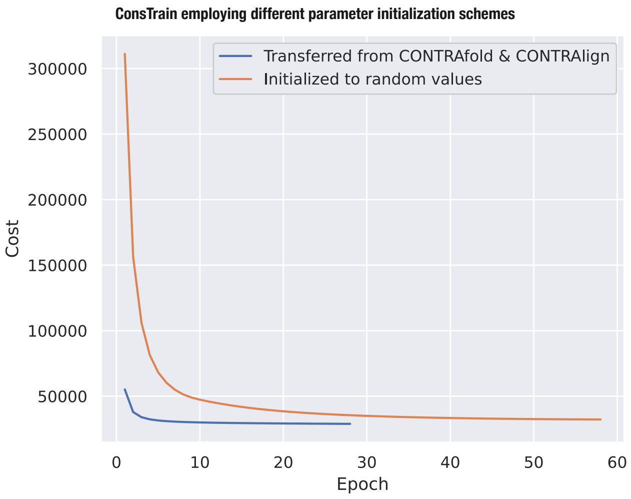


**Fig. S1.** Decrement of our L2-regularized non-convex cost in different parameter initialization settings. As the first choice, structural alignment scoring parameters were transfer-learned. As the second choice, structural alignment scoring parameters were initialized by random values. Dataset “train pairwise” was used.

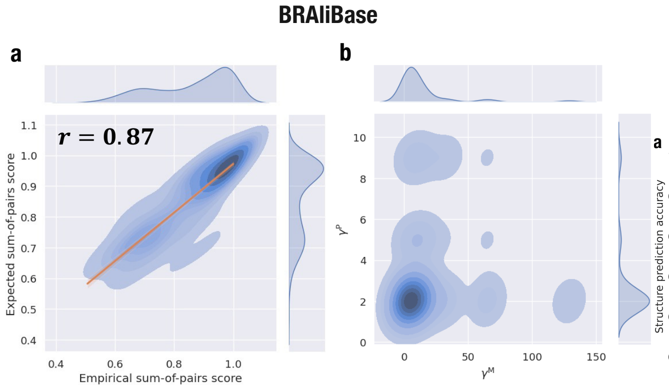


**Fig. S2.** The same as (a) Fig. 5 and (b) Fig. 6, except that BRAliBase's dataset 1 was used instead of the datasets "test multiple" and "RNAStralign processed."

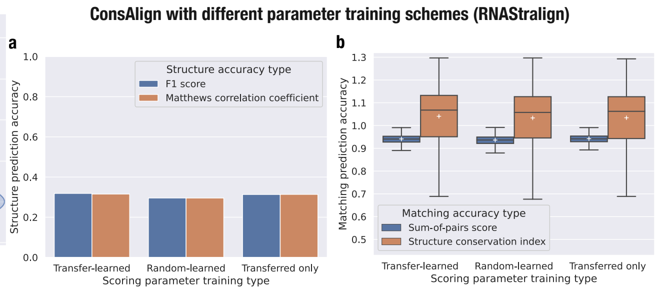


**Fig. S3.** The same as Fig. 7, except that the dataset "RNAStralign processed" was used instead of the dataset "test multiple."

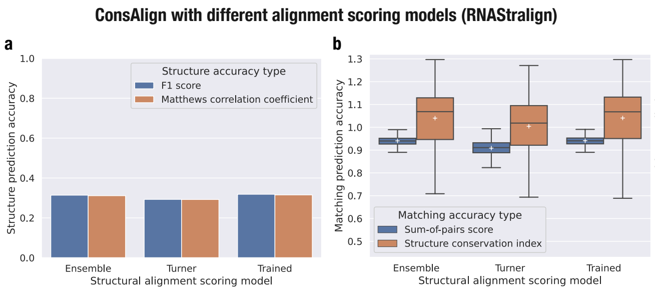


**Fig. S4.** The same as Fig. 8, except that the dataset "RNAStralign processed" was used instead of the dataset "test multiple."

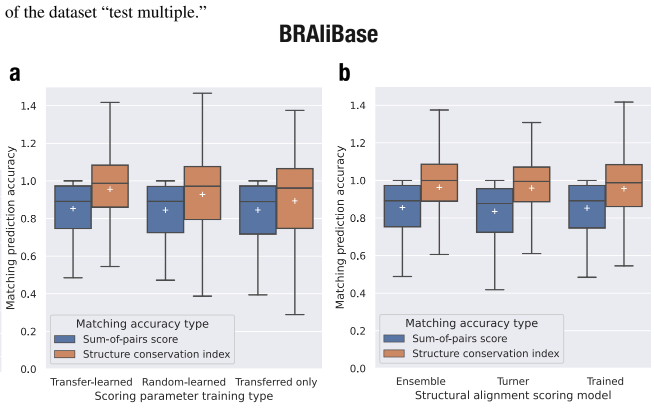


**Fig. S5.** The same as (a) Fig. 7b and (b) Fig. 8b, except that BRAliBase's dataset 1 was used instead of the dataset "test multiple."

### S4 Conventional structural aligners' features

(1) RAF (a) is a simultaneous RNA structural aligner taking both CONTRAfold and CONTRAlign's posterior probabilities and (b) trains the weight parameters of these posterior probabilities in max-margin optimization (Do *et al.*, 2008). (2, 3) LocARNA and SPARSE are different implementations of Sankoff's algorithm utilizing simultaneous structural alignment sparsity (Will *et al.*, 2007, 2015). SPARSE is a variant of LocARNA and exploits only structure-based constraints (Will *et al.*, 2015), whereas LocARNA exploits matching-based and structure-based constraints (Will *et al.*, 2007). (4) DAFS is a simultaneous RNA structural aligner and realizes reasonable computational complexities by applying dual decomposition to integer programming (Sato *et al.*, 2012). (5) LinearTurboFold is an application of LinearPartition (Zhang *et al.*, 2020) to TurboFold, an iterative RNA structural aligner (Tan *et al.*, 2017). LinearTurboFold realizes its quick predictive iteration using both LinearPartition and beam search-based sequence alignment (Li *et al.*, 2021).
